# Evolution of temperature dependence of TRPM2 channel gating and inactivation in vertebrates

**DOI:** 10.1101/2025.08.22.671705

**Authors:** Ádám Bartók, Ádám V. Tóth, László Csanády

## Abstract

TRPM2 is a Ca^2+^ permeable cation channel activated by cytosolic Ca^2+^ and ADP ribose (ADPR). In contrast to invertebrate orthologs, vertebrate TRPM2 channels inactivate due to evolutionary alterations in amino acid sequence around the selectivity filter. Human TRPM2 (hsTRPM2) serves as a deep-brain temperature sensor important for body temperature regulation, and its gating is exquisitely temperature dependent. To address whether TRPM2 is temperature sensitive also in ectotherms that lack body temperature regulation, here we investigated the functional properties of zebrafish (*Danio rerio*) TRPM2 (drTRPM2) across the temperature range of 15°C to 37°C. The rate of ion permeation through the open pore was weakly temperature sensitive (Q_10_∼1.3) as expected for a diffusion-limited process. In the presence of saturating concentrations of ligands (ADPR and Ca^2+^) drTRPM2 open probability showed no temperature dependence. Moreover, for both ADPR and Ca^2+^, the apparent affinities for channel activation were unaffected by temperature, reporting a standard enthalpy of opening near zero for drTRPM2. Inactivation of drTRPM2 is an order of magnitude slower than that of hsTRPM2. To address whether in both orthologs the same mechanism underlies inactivation, we studied its temperature sensitivity. For both drTRPM2 and hsTRPM2 inactivation rate was modestly temperature dependent (Q_10_ ∼2.7 and ∼4.4). A triple substitution which converts the post-filter sequence of drTRPM2 into the corresponding human sequence accelerated drTRPM2 inactivation by >20-fold. The data suggest that temperature dependence of hsTRPM2 gating evolved in the course of vertebrate evolution, whereas inactivation was temperature dependent already when it first appeared in early vertebrates.

## Introduction

Transient Receptor Potential Melastatin 2 (TRPM2) is a Ca^2+^-permeable nonselective cation channel activated by binding of cytosolic ADP ribose (ADPR) and Ca^2+^ [1]. In mammals TRPM2 plays important roles in immune cell activation [2], insulin secretion [3]), and the central regulation of body temperature [4, 5].

The pore-forming transmembrane domain (TMD) of a homotetrameric TRPM2 channel (structural Layer 1) is formed from six transmembrane helices (S1-S6), and its architecture resembles that of voltage-gated cation channels [6, 7]. Within the long extracellular loop that links S5 to S6, a short pore helix followed by a pore loop forms the selectivity filter. The channel gate is lined by the C-termini of S6 which form a tetrahelix bundle at the cytosolic membrane surface. In the immediate vicinity of the channel gate, the S2-S3 linkers form highly conserved binding sites for activating Ca^2+^ ions [6-8]. Below the TMD, large N- and C-terminal cytosolic regions intertwine to build a three-layered structure (Layers 2-4) which houses two binding sites per subunit for ADPR. A first binding site in Layer 3, formed by an N-terminal domain (N-site) is primarily responsible for channel gating [7-9]. A second binding site in Layer 4, formed by the C-terminal NUDT9-homology (NUDT9H) domain (C-site), has adopted different functions in different orthologs: in invertebrate TRPM2 channels it is an active ADPR hydrolase (ADPRase) but plays no role in channel gating [10, 11], in vertebrate orthologs it is enzymatically inactive and its role in channel gating is unclear [12]. The structural organization of Layers 1-3 has remained largely conserved from choanoflagellates to man [6-8, 13].

For all TRPM2 orthologs, channel opening requires simultaneous binding of Ca^2+^ and ADPR, as well as the presence of membrane phosphatidylinositol-4,5-bisphosphate (PIP_2_) [10, 14, 15]. However, whereas invertebrate TRPM2 channels remain active as long as cytosolic ligands are supplied [10], the human channel (hsTRPM2) irreversibly inactivates when kept open by the prolonged presence of agonists [14]. The evolutionary appearance of that inactivation coincided with sequence alterations, between invertebrates and vertebrates, of a specific amino acid triplet in the short helix that follows the selectivity filter (‘post-filter helix’): a single-residue deletion and the loss of two negative charges [10]. Replacement of that triplet in the sea anemone (*Nematostella vectensis*) ortholog (nvTRPM2) with the corresponding human doublet induces inactivation in nvTRPM2 [6, 10], whereas introduction of the nvTRPM2 triplet into hsTRPM2 eliminates inactivation [6, 15]. Inactivation of vertebrate TRPM2 channels thus likely reflects a conformational change of the selectivity filter. Interestingly, for the zebrafish (*Danio rerio*) ortholog (drTRPM2), one of the most ancient vertebrate TRPM2 channels, the rate of inactivation is an order of magnitude slower than for hsTRPM2 [10].

TRPM2 belongs to the subset of TRP channels called ‘thermoTRPs’ for their involvement in temperature sensation [16]. In mammals TRPM2 is expressed in neurons of the thermoregulatory center of the brain, the preoptic area (POA) of the hypothalamus. TRPM2 activity in the POA reports deep-brain temperature, and strongly impacts thermoregulation [4, 5]. To achieve that aim, TRPM2 activity must respond to the tiny temperature fluctuations of the brain, typically as small as ±1°C. Correspondingly, biophysical studies on hsTRPM2 have found its activity to be exquisitely temperature sensitive: intrinsic temperature-dependence of gating is greatly enhanced by a positive feedback mechanism caused by Ca^2+^ influx through the open pore. As a result, hsTRPM2 open probability increases by ∼2-fold per 1°C (Q_10_∼1000) at physiological ligand concentrations and temperature (∼37°C) [17].

An interesting question is whether temperature-sensitivity was a pre-existing biophysical property of TRPM2 which became exploited in endotherms to employ TRPM2 as a deep-brain thermometer, or whether temperature sensitivity of TRPM2 gating evolved together with the appearance of body temperature regulation. Moreover, although inactivation is also important for shaping TRPM2 activity time courses, its temperature dependence has not yet been studied for any ortholog. We therefore undertook a detailed study of temperature sensitivity of gating and inactivation for TRPM2 from a representative ectothermic early vertebrate. We chose to study drTRPM2 for which a large body of structural and functional information is already available. Previous reports have assessed the properties of other thermoTRPs, such as TRPV1 [18] and TRPA1 [19] expressed in the zebrafish, but for drTRPM2 such detailed studies have not yet been published.

## Materials and Methods

### Molecular biology

The drTRPM2/pcDNA3.0 construct was generated from drTRPM2/pGEMHE [10] by subcloning the coding region into pcDNA3.0. The hum-drTRPM2/pcDNA3.0 construct was generated from drTRPM2/pcDNA3.0 using the QuikChange II XL Site-Directed Mutagenesis Kit (Agilent Technologies). Primers (Forward: CTTCGGCAACATTCCCGGTTACATTGACAACACCCTGTTCG; Reverse: CGAACAGGGTGTTGTCAATGTAACCGGGAATGTTGCCGAAG) were purchased from Sigma-Aldrich. The construct was confirmed by automated sequencing (LGC Genomics).

### Expression of TRPM2 constructs in HEK-293 cells

HEK 293T cells were obtained from ATCC (CRL-11268) and cotransfected with drTRPM2/pcDNA3.0 (or hum-drTRPM2/pcDNA3.0) and GFP/pcDNA3 at a 10:1 ratio (FuGENE HD transfection reagent, Promega). HEK 293 cells stably expressing hsTRPM2 were purchased from SB Drug Discovery. All cells were cultured at 37°C in 5% CO_2_ in DMEM medium supplemented with 4.5 g/L Glucose (Lonza), 10% FBS (EuroClone), 2 mM glutamate, and 100 units/ml penicillin/streptomycin (Lonza).

### Inside-out patch-clamp recordings

Currents from drTRPM2 (hum-drTRPM2) or hsTRPM2 channels were recorded in inside-out patches from transiently or stably expressing HEK 293 cells, respectively, as described [17]. Pipette (extracellular) solution contained (in mM) 140 Na-gluconate, 2 Mg-gluconate_2_, 10 PIPES (pH 7.4 with NaOH at 25°C) with or without 1 EGTA (pH 7.4 with NaOH at 25°C). A 140 mM NaCl-based solution for the pipette electrode was carefully layered on top [14]. Bath (cytosolic) solution contained (in mM) 140 Na-gluconate, 2 Mg-gluconate_2_, 10 PIPES (pH 7.1 with NaOH). Free [Ca^2+^] in the micromolar range was adjusted by titrating gluconate with Ca^2+^. For solutions with 1 nM - 1 μM free [Ca^2+^] 1 mM EGTA was added and titrated with Ca^2+^. Free [Ca^2+^] was spectrophotometrically determined at each temperature [17]. Temperature-dependent changes in pH of the pipette and bath solutions were kept minimal by the use of PIPES buffer (ΔpK_a_/10°C=-0.085). All bath solutions also contained 200 μM AMP (to block endogenous TRPM4-like cation channels) and 10 μM dioctanoyl-PIP_2_. The bath electrode (in 3 M KCl) was connected to the cytosolic solution through a KCl-agar bridge. The continuously flowing bath solution was exchanged (time constant <100 ms) using electronic valves (ALA-VM8, ALA Scientific Instruments). Flow line temperatures were controlled (TC-10 Dagan Corporation), and bath temperature was continuously monitored (BAT-12, Physitemp) at a distance of ∼2 mm from the membrane patch [20]. Ca^2+^ (Ca-gluconate_2_; Sigma-Aldrich), ADPR (Sigma-Aldrich), dioctanoyl-PIP_2_ (Cayman Chemical), and AMP (Sigma-Aldrich) were dissolved into the bath solution from >100x concentrated, pH-adjusted aqueous stocks. Currents were recorded at a bandwidth of 2 kHz (Axopatch 200B; Molecular Devices), digitized at 10 kHz (Digidata 1322A; Molecular Devices), and saved to disk (Pclamp10; Molecular Devices)

### Data analysis

Macroscopic fractional currents (Figs. 1D, 2D, 2H) were calculated as the mean steady-state current under a test condition (i.e., test concentration of an agonist, or test temperature) normalized to the average of the mean steady-state currents observed in the same patch in bracketing segments of record under reference conditions (i.e., in the presence of maximal agonist concentrations, or at the reference temperature (25°C)). Dose response curves were fitted to the Hill equation using least squares. Macroscopic current decay time courses were fitted using least-squares to decaying single-exponential functions to obtain time constants (Figs. 3B, 3D, and 4C).

**Figure 1.**
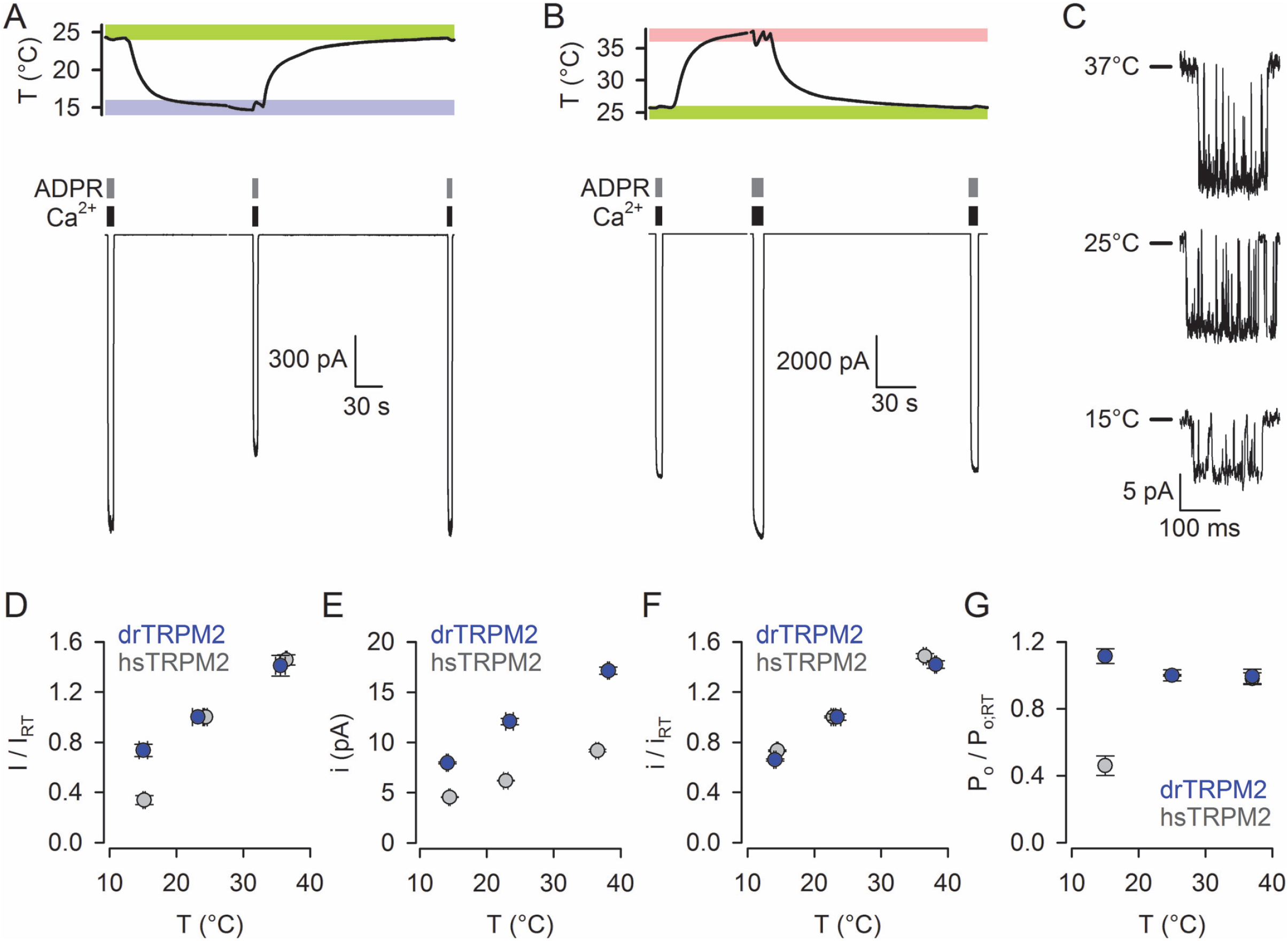
Gating of fully liganded drTRPM2 channels is insensitive to temperature between 15°C and 37°C. *A*-*B, Bottom*, Macroscopic drTRPM2 currents in inside-out patches from HEK-293T cells evoked by brief exposures to saturating concentrations of cytosolic Ca^2+^ (111-148 μM; *black bars*) and ADPR (32 μM; *gray bars*) at temperatures alternating between ∼25°C and ∼15°C (*A*) or between ∼25°C and ∼37°C (*B*). Membrane potential was -80 mV. *Top*, Bath temperatures recorded in the immediate vicinity of the patch. *Blue, green*, and *rose boxes* highlight target temperature ranges of 15±1°C, 25±1°C, and 37±1°C, respectively. *C*, Segments of unitary drTRPM2 current at -80 mV at the three target temperatures. *D*-*G*, Temperature dependence of normalized macroscopic current (*D*), absolute unitary current amplitude (*E*), normalized unitary current amplitude (*F*), and normalized P_o_ (*G*) for drTRPM2 (*blue symbols*) and hsTRPM2 (*gray symbols*; replotted from [17]). The parameters in panels *D, F*, and *G* were normalized to their mean values at ∼25°C (‘RT’). Symbols represent mean±S.E.M. For drTRPM2 n=4-6 (*D*) and n=19-36 (*E*-*F*). For hsTRPM2 n=8-26 (*D*) and n=4-6 (*E*-*F*). In panel *G*, S.E.M. was calculated from the variances of I/I_RT_ and i/i_RT_ by error propagation.

### Estimation of unitary current amplitudes

Current amplitude histograms were fitted by sums of Gaussian functions, and unitary current amplitudes (Fig. 1C) estimated as the distances between adjacent peaks.

### Statistics

All data are shown as mean±S.E.M., with the number of independent experiments (n) indicated in the figure legends.

## Results

### Gating of fully liganded drTRPM2 channels is insensitive to temperature between 15°C and 37°C

To study its biophysical properties, drTRPM2 was expressed in HEK-293T cells and studied in excised inside-out patches at -80 mV membrane potential, in symmetrical Na-gluconate based solutions. Macroscopic TRPM2 currents (Fig. 1A-B, *Bottom*) and bath temperature in the immediate vicinity of the patch (Fig. 1A-B, *Top*) were simultaneously recorded. Continuous direct superfusion of the cytoplasmic patch surface ensured rapid addition/removal of ligands (Fig. 1A-B, *Bottom, black* and *gray bars*), and a temperature controller afforded ramping the temperature between our control temperature of ∼25°C (Fig. 1A-B, *Top, green boxes*) and either ∼15°C (Fig. 1A, *Top, blue box*) or ∼37°C (Fig. 1B, *Top, rose box*) (Materials and Methods; [17]).

The slow rate of drTRPM2 inactivation (time constant >100 s at 25°C; [10]) allowed estimation of macroscopic TRPM2 currents by brief (5-10 s) exposures to saturating concentrations of both Ca^2+^ (111-148 μM, see Materials and Methods) and ADPR (32 μM), without causing substantial inactivation (Fig. 1A-B, Bottom). Ligand-activated currents evoked at the test temperatures of ∼15°C (Fig. 1A) or ∼37°C (Fig. 1B) were normalized to the averages of the currents evoked in the same patch in bracketing segments of record at the control temperature of ∼25°C (‘Room Temperature’, ‘RT’). The plot of normalized macroscopic current (*I*/*I*_RT_) vs. temperature (Fig. 1D, *blue symbols*) was roughly linear and showed mild temperature dependence. To dissect the contributions of changes in open probability (P_o_) and unitary current amplitude (*i*) to overall temperature sensitivity of drTRPM2 currents, unitary amplitudes (Fig. 1C) were also quantitated and plotted against temperature, yielding a linear *i*(T) plot (Fig. 1E, *blue symbols*). Although the unitary conductance of drTRPM2 is almost twice as large as that of hsTRPM2 (Fig. 1E, *gray symbols*, replotted from [17]), the fractional effect of temperature, obtained by normalizing unitary currents to those at ∼25°C (*i*/*i*_RT_) was essentially identical for both orthologs (Fig. 1F, *blue* vs. *gray symbols*), and consistent with temperature dependence of the rate of diffusion of ions in free solution (Q_10_∼1.3). The macroscopic current is the product of the unitary current, the open probability, and the number of channels in the patch (*I* =*i*·P_o_·*N*). Thus, the fractional effect of temperature on open probability (P_o_/P_o;RT_) is obtained as P_o_/P_o;RT_ = (*I*/*I*_RT_) /(*i*/*i*_RT_). Whereas for hsTRPM2 that analysis revealed a marked decline in P_o_ upon cooling from 25°C to 15°C (Fig. 1G, *gray symbols*; replotted from [17]), for drTRPM2 (Fig. 1G, *blue symbols*) no decline in P_o_ was detectable. Thus, in contrast to hsTRPM2, for drTRPM2 gating at saturating ligand concentrations shows no temperature dependence between 15°C and 37°C.

### Apparent affinities of drTRPM2 for ADPR and Ca^2+^ are unaffected by temperature between 15°C and 37°C

The apparent lack of temperature dependence of P_o_ of fully liganded drTRPM2 within the limited temperature span tested here may either reflect a small standard enthalpy of opening (ΔH^0^), or may alternatively be caused by a temperature threshold which falls far below the investigated range of temperatures. However, in the latter case temperature dependence should appear at lower, subsaturating ligand concentrations, manifested by rightward shifts in ligand dose-response curves at lower temperatures. E.g., for hsTRPM2 ΔH^0^ is large (∼180 kJ/mol), but in the presence of saturating ligands the temperature threshold is <15°C [17], so that P_o_ is essentially saturated and shows little temperature dependence at T>25°C (cf., Fig. 1G, *gray symbols*). On the other hand, apparent affinities for both ligands decrease by ∼5-fold from 40°C to 25°C [17].

To differentiate between those two possibilities, ligand concentration dependences of macroscopic current activation were assessed at ∼15°C (Fig. 2A, E), ∼25°C (Fig. 2B, F), and ∼37°C (Fig. 2C, G) for both ADPR (Fig. 2A-C) and Ca^2+^ (Fig. 2E-G), by normalizing steady-state currents during brief application of a test ligand concentration to the averages of the currents during bracketed applications of a saturating concentration, while keeping the other ligand saturating. The resulting dose response curves (Fig. 2D, H; *colored symbols*) were fitted by the Hill equation to obtain apparent K_1/2_ values (Fig. 2D, H; *colored curves* and *numbers*). Intriguingly, the dose response curves obtained at the three temperatures essentially overlay with each other for both ADPR and Ca^2+^ – in stark contrast to the large temperature-dependent shifts in K_1/2_ reported earlier for hsTRPM2 [17]. These results suggest a virtual lack of temperature dependence, i.e., a small ΔH^0^, for the closed-open conformational transition of drTRPM2.

**Figure 2.**
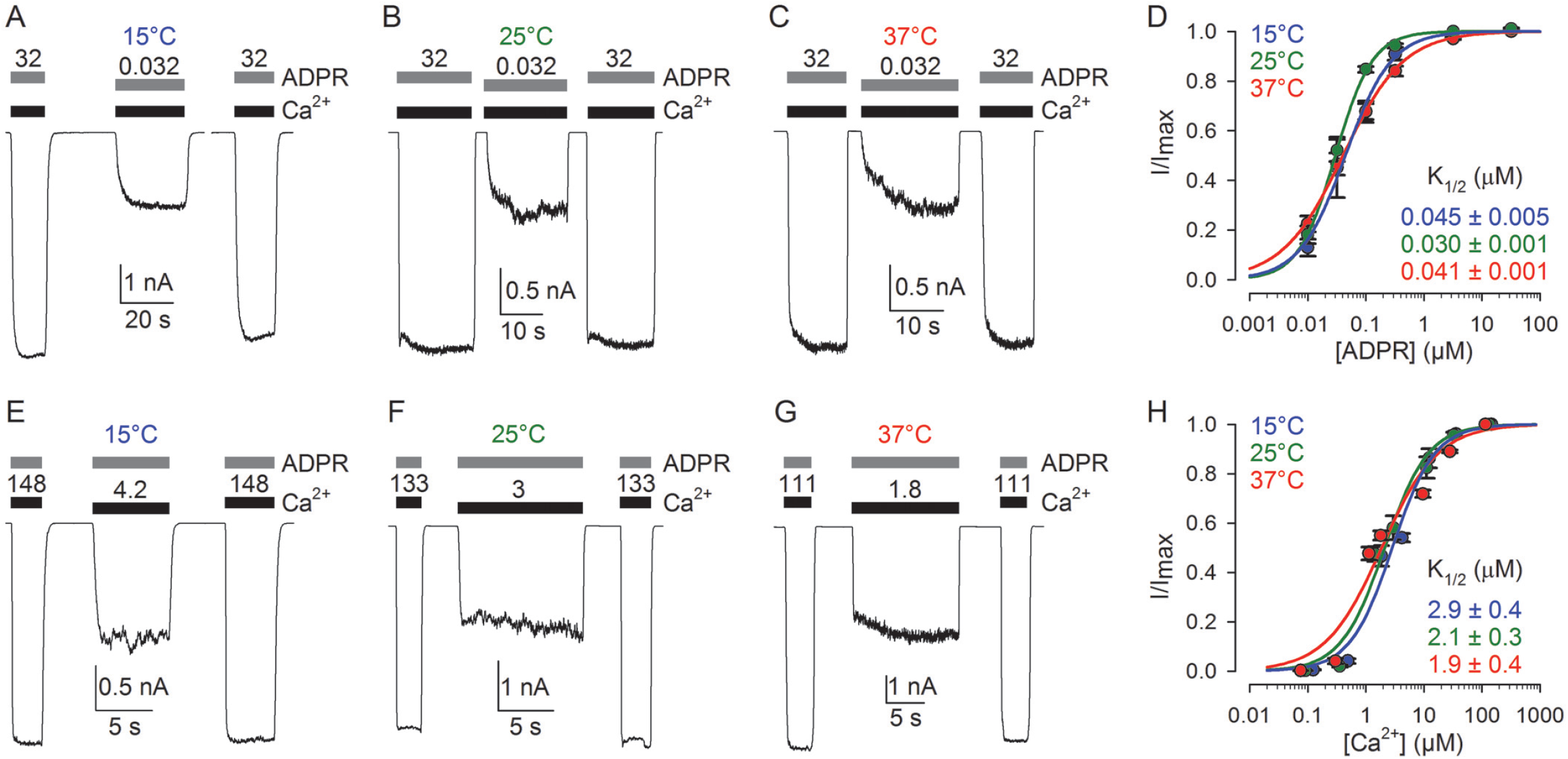
Ligand dose response curves of drTRPM2 are unaffected by temperature. *A*-*C, E*-*G*, Macroscopic drTRPM2 currents evoked in inside-out patches by alternating brief exposures to saturating and subsaturating concentrations of either ADPR (*A*-*C*) or Ca^2+^ (*E*-*G*), together with a saturating concentration of the other ligand. Bath temperature was ∼15°C (*A, E*), ∼25°C (*B, F*), or ∼37°C (*C, G*). Membrane potential was -80 mV. *D, H*, Normalized dose response curves for (*D*) ADPR in the presence of saturating Ca^2+^, and (*H*) for Ca^2+^ in the presence of saturating ADPR, at the three target temperatures (*color coded*). Fractional current in the presence of a test ligand concentration was normalized to the average of the currents evoked in the same patch by bracketed exposures to a saturating ligand concentration. Symbols plot mean±S.E.M. In *D*, n=8-15 (15°C), n=5-13 (25°C), n=12-21 (37°C). In *H*, n=4-9 (15°C), n=4-8 (25°C), n=4-11 (37°C). *Solid curves* are fits to the Hill equation with midpoints (K_1/2_) plotted in the panels.

### Inactivation rate is modestly temperature dependent for both drTRPM2 and hsTRPM2

Temperature dependence of TRPM2 inactivation has so far not been systematically addressed. We therefore studied the rates of inactivation for both drTRPM2 (Fig. 3A) and hsTRPM2 (Fig. 3C), by recording macroscopic currents in the prolonged presence of saturating concentrations of both ADPR and Ca^2+^, at 37°C (Fig. 3A, C; *red traces*) and 25°C (Fig. 3A, C; *green traces*). Inactivation time constants (Fig. 3B, D), obtained by single-exponential fits to the current decay time courses, showed significant temperature dependence for both orthologs: compared to 37°C, the inactivation time constant was prolonged by ∼3.2-fold for drTRPM2 (Q_10_∼2.7), and by ∼6.0-fold for hsTRPM2 (Q_10_∼4.4). Thus, inactivation was temperature dependent already when it first appeared in early vertebrates suggesting that – despite the difference in absolute rates between the two orthologs – its mechanism is similar in drTRPM2 and hsTRPM2.

**Figure 3.**
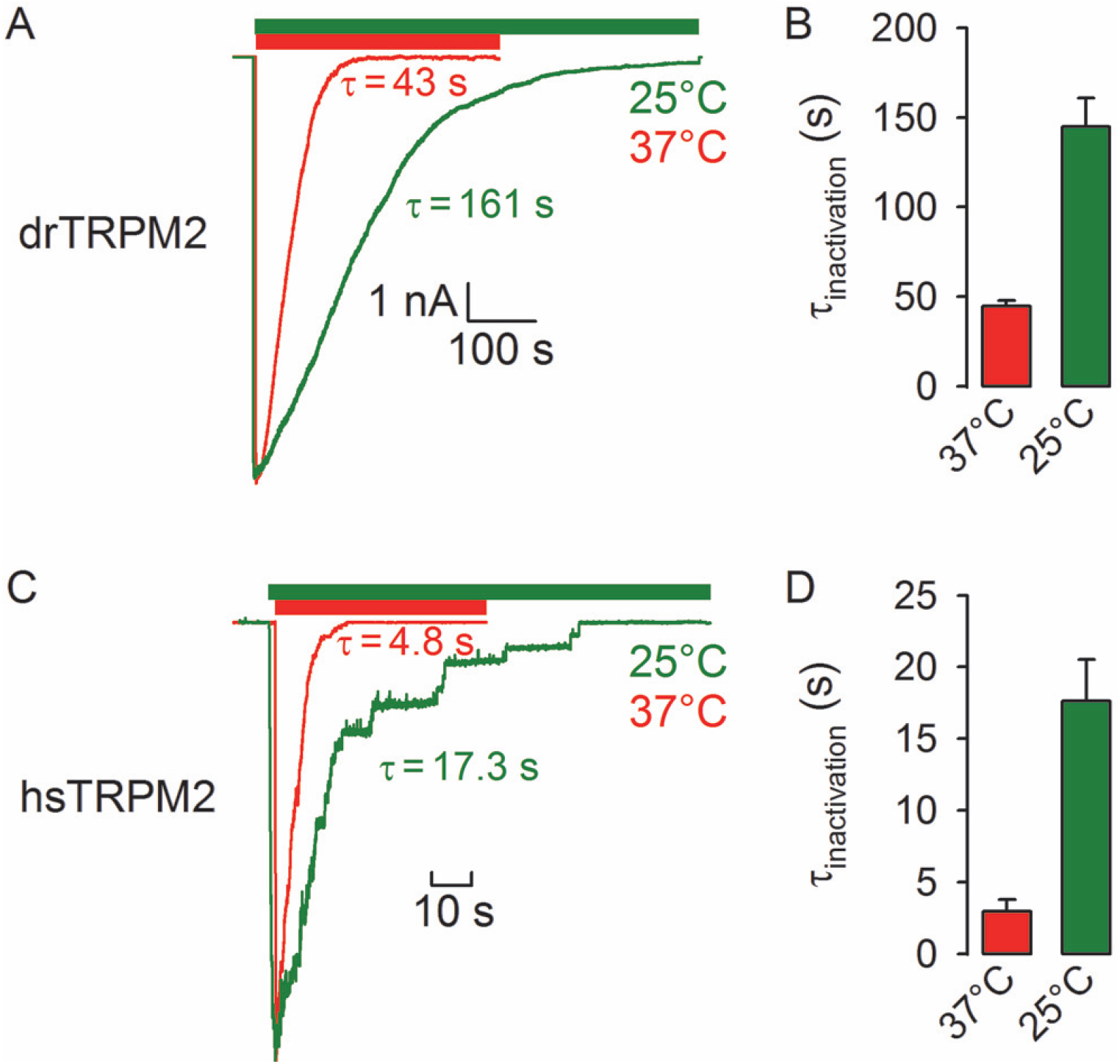
Inactivation rate is temperature dependent for both drTRPM2 and hsTRPM2. *A, C*, Macroscopic currents of drTRPM2 (*A*) and hsTRPM2 (*C*) channels in inside-out patches evoked by sustained exposure (*colored bars*) to saturating Ca^2+^ (133-148 μM) plus ADPR (32 μM) at ∼25°C (*green traces* and *bars*) or ∼37°C (*red traces* and *bars*). Membrane potential was -80 mV. In *C*, the current traces were rescaled by their maximal amplitudes. *B, D*, Time constants of inactivation for drTRPM2 (*B*) and hsTRPM2 (*D*) at ∼37°C (*red*) and ∼25°C (*green*), obtained from single-exponential fits to the traces in *A, C*. Data represent mean±S.E.M. In *B*, n=8 (25°C), n=39 (37°C). In *D*, n=9 (25°C), n=7 (37°C).

### drTRPM2 with a ‘humanized’ post-filter sequence inactivates as fast as hsTRPM2

Following the single-residue deletion and the loss of two negative charges from the post-filter helix, which happened between invertebrates and early vertebrates (Fig. 4A, compare *1st* and *2nd row*), three additional substitutions in the post-filter region took place between zebrafish and humans (Fig. 4A, compare *2nd* and *3rd row*). To address whether the latter substitutions are responsible for the largely accelerated rate of inactivation of hsTRPM2 relative to drTRPM2, a ‘humanized’ drTRPM2 construct (‘hum-drTRPM2’) was generated by introducing the same three substituions into drTRPM2 (Fig. 4A, *4th row*). Indeed, inactivation rate of hum-drTRPM2 (Fig. 4B) was accelerated by ∼20-fold relative to drTRPM2 (Fig. 4C), rendering it comparable to the inactivation rate of hsTRPM2.

**Figure 4.**
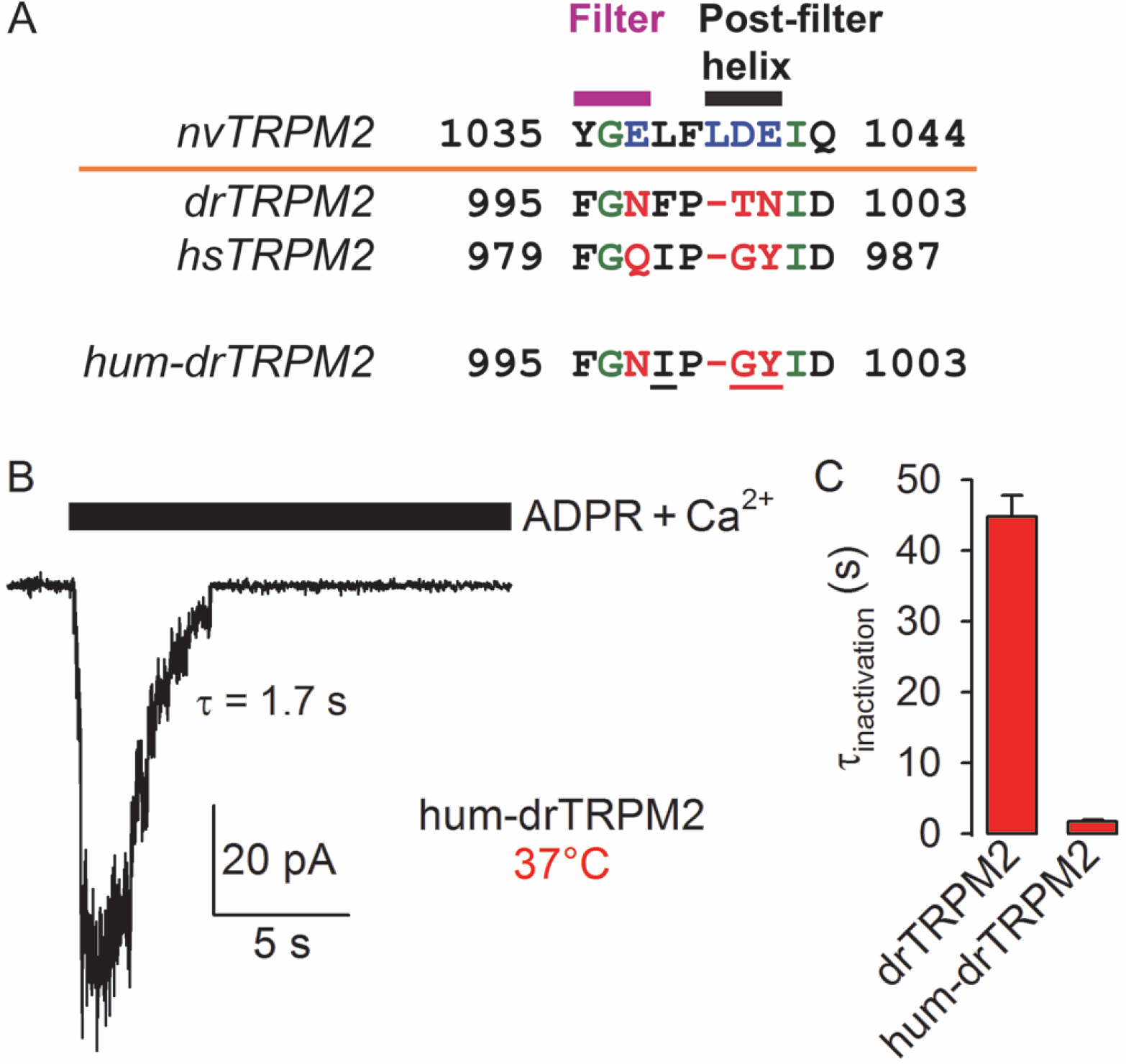
‘Humanizing’ the drTRPM2 post-filter sequence accelerates inactivation. *A*, Sequence alignment of the filter/post-filter region for nvTRPM2, drTRPM2, and hsTRPM2, as well as for the hum-drTRPM2 construct. Residues mutated in hum-drTRPM2 are underlined. *B*, Macroscopic current of hum-drTRPM2 channels in an inside-out patch evoked by sustained exposure (*black bar*) to saturating Ca^2+^ (133 μM) plus ADPR (32 μM) at ∼37°C. Membrane potential was -80 mV. *C*, Time constants of inactivation of drTRPM2 (*left*, replotted from Fig. 3B) and hum-drTRPM2 (*right*) at ∼37°C. Data represent mean±S.E.M. For hum-drTRPM2, n=9.

## Discussion

Zebrafish body temperature adopts its environmental temperature, which ranges from ∼10 °C to ∼40 °C in various natural habitats of this species [21]. In the present study we compared functional properties (conductance, gating, and inactivation) of drTRPM2 across most of that temperature range.

We found that gating of drTRPM2 lacks temperature dependence (ΔH^0^∼0), in contrast to the strong intrinsic temperature dependence of hsTRPM2 gating (ΔH^0^∼180 kJ/mol; [17]). This finding is intriguing, given that the gating mechanisms of the two orthologs are otherwise very similar, both being activated by ligand binding to structurally highly conserved binding sites: Ca^2+^ binding to the S2-S3 linker and high-affinity ADPR binding to the N-site [7, 8, 12, 22-24]. Moreover, even the conformational rearrangements observed upon ligand binding are similar, at least for structural Layers 1-3 [7, 8]. We show here that for drTRPM2 apparent K_1/2_ values for both ligands are tuned to be similar to those reported for hsTRPM2 at 40°C [17], ensuring high-affinity activation of the zebrafish channel across the entire temperature range encountered by this species.

From a biophysical point of view, an interesting question is which structural parts of hsTRPM2 are responsible for its temperature sensitivity. In that regard, the various thermoTRP channels appear to employ diverse sets of mechanisms. For the close homolog TRPM4 which shares, from a functional point of view, Ca^2+^-activation and, from a structural point of view, Layers 1-3 with TRPM2, temperature sensitivity has been proposed to result from a ‘temperature-sensitive Ca^2+^ binding site’ in the N-terminal cytosolic region, distinct from the conventional Ca^2+^ site formed by the TMDs [25]. However, because the Ca^2+^-coordinating side chains of that cytosolic Ca^2+^ site are not conserved in TRPM2, it seems unlikely that the mechanism proposed to underly TRPM4 temperature dependence would also apply to TRPM2. One structural difference between drTRPM2 and hsTRPM2 is the arrangement of Layer 4 in the unliganded state. Due to intersubunit interactions between the N-terminal region and a short segment (the ‘P-loop’) of the unliganded NUDT9H domain, unliganded Layer 4 forms a closed gating ring in hsTRPM2, but not in drTRPM2 which lacks a P-loop [7, 8]. Indeed, it has been shown that the NUDT9H P-loop of hsTRPM2 does play a role in its temperature sensitivity [26]. However, to confound the picture, the P-loop is also present in TRPM2 orthologs of other ectothermic vertebrates (amphibians and reptiles) and even of more ancient species such as cartilaginous fishes and invertebrates, suggesting that it was selectively deleted in bony fishes. Thus, detailed biophysical studies on the structural organization and temperature dependence of more TRPM2 orthologs will be required to shed light on the exact structural underpinnings of hsTRPM2 temperature sensitivity.

TRPM2 inactivation, which appeared in early vertebrates, reflects pore instability due to a triple substitution in the outer pore ([6, 10, 15], Fig. 4A). Here we show that further acceleration of that process in hsTRPM2 is caused by additional substitutions in the same region (Fig. 4B). The modest temperature dependence of that process is unlikely to be physiologically relevant for TRPM2 expressed in the human brain, where the temperature is maintained within a narrow range of 1-3°C. In contrast, it might potentially play a physiological role in zebrafish, the body temperature of which adopts a large range (>30°C) of environmental temperatures.

Zebrafish are responsive to environmental temperature gradients [18], which implies the ability to sense temperature. From a physiological point of view, a question of interest is which thermoTRP channels contribute to its temperature sensing. Such a role has already been demonstrated for drTRPV1 [18] and for one of two drTRPA1 paralogs [19, 27]. Future studies will need to address whether drTRPM2, which is expressed in various populations of sensory neurons during different stages of zebrafish development [28], contributes to temperature sensing at all. In principle, the temperature dependence of its inactivation might become relevant at extreme environmental temperatures.

## Acknowledgments

Supported by EU Horizon 2020 Research and Innovation Program grant 739593, and National Research, Development and Innovation Fund grants KKP 144199 to L.C. and ADVANCED 149640 to A.B. A.B. was supported by the János Bolyai Research Scholarship of the Hungarian Academy of Sciences (BO/00103/20 and BO/00238/25/8) and the New National Excellence Program (ÚNKP) Bolyai+ scholarship of the Ministry of Human Capacities of Hungary (ÚNKP-20-5-SE-6, ÚNKP-21-5- SE-10 and ÚNKP-22-5-SE-12). A.V.T. was supported by the 2024-2.1.1-EKÖР-2024-00004 University Research Scholarship Programme of the Ministry for Culture and Innovation of Hungary (source National Research, Development and Innovation Fund).

## Data availability statement

All data generated over the course of this study are included within the paper.

## CRediT authorship contribution statement

A.B.: Conceptualization, Data curation, Formal analysis, Funding acquisition, Investigation, Methodology, Visualization, Writing – original draft. A.V.T.: Funding acquisition, Investigation. L.C.: Conceptualization, Data curation, Formal analysis, Funding acquisition, Methodology, Visualization, Writing – original draft.

